# An intrinsically interpretable neural network architecture for sequence to function learning

**DOI:** 10.1101/2023.01.25.525572

**Authors:** Ali Tuğrul Balcı, Mark Maher Ebeid, Panayiotis V Benos, Dennis Kostka, Maria Chikina

## Abstract

**Motivation:** Sequence-based deep learning approaches have been shown to predict a multitude of functional genomic readouts, including regions of open chromatin and RNA expression of genes. However, a major limitation of current methods is that model interpretation relies on computationally demanding post hoc analyses, and even then, one can often not explain the internal mechanics of highly parameterized models. Here, we introduce a deep learning architecture called tiSFM (totally interpretable sequence to function model). tiSFM improves upon the performance of standard multi-layer convolutional models while using fewer parameters. Additionally, while tiSFM is itself technically a multi-layer neural network, internal model parameters are intrinsically interpretable in terms of relevant sequence motifs.

**Results:** We analyze published open chromatin measurements across hematopoietic lineage cell-types and demonstrate that tiSFM outperforms a state-of-the-art convolutional neural network model custom-tailored to this dataset. We also show that it correctly identifies context specific activities of transcription factors with known roles in hematopoietic differentiation, including Pax5 and Ebf1 for B-cells, and Rorc for innate lymphoid cells. tiSFM’s model parameters have biologically meaningful interpretations, and we show the utility of our approach on a complex task of predicting the change in epigenetic state as a function of developmental transition.

**Availability and implementation:** The source code, including scripts for the analysis of key findings, can be found at https://github.com/boooooogey/ATAConv, implemented in Python.

## 1 Introduction

Functional genomic assays have accelerated our understanding of how non-coding regions of genomes contribute to both cellular and organismal function through their identification and characterization. Large consortium projects such as ENCODE (ENCODE Project Consortium, 2012; Gal-Oz *et al*., 2019) have generated extensive datasets profiling diverse functional properties of the non-coding genome across many tissues and cell-types, including transcription factor (TF) binding, regions of open chromatin, and biochemical modification of N-terminal histone tails. While such genomic readouts are believed to be largely determined by DNA sequence, the precise sequence-to-function (S2F) relation is complex and remains poorly understood. Nevertheless, recent developments have shown that by using deep learning models, with millions of parameters, it is indeed possible to learn S2F mappings, predict epigenetic readouts, or even characterize gene expression (Maslova *et al*., 2020; Avsec *et al*., 2021b; Kelley *et al*., 2016; Zhou and Troyanskaya, 2015; Quang and Xie, 2016; Dibaeinia and Sinha, 2021; Avsec *et al*., 2021a). Given these successes, it is worthwhile to ask what scientific insights that go beyond advancing our understanding of applying machine learning engineering principles to genomics data such models can provide.

Primarily, a model can be considered useful (in the canonical, supervised machine learning sense) if it is able to make reliable out-of-sample predictions. This is a major promise of complex deep learning models, given they can potentially predict functional output of arbitrary sequences (e.g., individual genomes, haplotypes with disease-associated genetic variants, or synthetic constructs). However, while existing models have made extensive progress toward this goal, existing state-of-the-art models are not yet capable of supplanting direct measurement (Zhou and Troyanskaya, 2015; Avsec *et al*., 2021a).

Moving beyond out-of-sample predictions, a successful model can also provide information about the underlying biochemical principles, mechanisms, or interactions. For example, if increasing the receptive field of a S2F model from 40 kb to 200 kb increases model performance (as was recently demonstrated for gene expression predictions in Avsec *et al*., 2021b), then we can conclude that there is indeed evidence for relevant biochemical interactions that occur in the scope of 200 kb. However, questions about the actual biochemical entities involved, or how they may interact, or be organized into higher order structures, cannot be answered from observing model performance. Instead, model parameters need to be amendable to transparent post hoc interpretation. The architectures of many deep learning S2F models include an initial convolution layer whose kernels can be interpreted as position weight matrices (PWMs), and matched to databases of known transcription factor motifs, thereby revealing key regulators (Alipanahi *et al*., 2015; Quang and Xie, 2016; Liu *et al*., 2020; Maslova *et al*., 2020).

Establishing these overall goals emphasizes that the successful incorporation of such strategies is highly dependent on a model’s hyperparameters. From the beginning, learned kernel weights may only represent partial TF motifs that are aggregated in subsequent layers of the network (Zhou and Troyanskaya, 2015; Quang and Xie, 2016; Banovich *et al*., 2017; Koo and Eddy, 2019; Liu *et al*., 2020). Moreover, even if this aggregation in subsequent layers is successful, since neither the sign nor the magnitude of a TF’s contribution is known, kernel analysis approaches can only reveal which TFs are involved, but not how they contribute. These specific contributions can be evaluated through attribution analyses (a class of algorithms that can attribute a prediction to a specific subset of the input), highlighting regions of input DNA that can subsequently be matched to motifs of known TFs. This overcomes the problem of partial motifs, allowing for specific contributions to be evaluated. However, attribution analyses cannot provide a mechanistic interpretation for internal layers and require considerable computation and expertise (Dibaeinia and Sinha, 2021), as the results will vary depending on the method chosen.

The existing work that has combined kernel interpretation with attribution has led to significant progress in revealing both the transcription factors (TFs) involved and their specific contributions (sign and magnitude) towards predicting functional genomic readouts, as demonstrated by `AI-TAC` (Maslova *et al*., 2020). Specifically, `AI-TAC` focused on a hematopoeitic development dataset, and combined attribution and kernel analysis to create a TF by cell-type map. Despite being effective in identifying many known drivers of hematopoietic differentiation, `AI-TAC` required a synthesis of complex post-processing steps. Additionally, while the interpretation of the product, the TF by cell-type contribution, partially reveals the underlying mechanism of prediction performance, it does not provide insights into how the internal parameters of the model contribute to the predictions. For example, the `AI-TAC` method has 3 convolutions layers and 3 linear layers, an architecture that was extensively optimized for the dataset. However, the kernel-motif matching combined with attribution tells us nothing about how the internal parameters of the model contribute to the predictions. Do subsequent convolution layers encode interactions or simply fine-tune motifs? When multiple motifs contribute to a prediction does the model learn and OR, AND, or simply additive function?

In this work, we introduce a “totally interpretable sequence-to-function model”, `tiSFM`. While the model is technically a multi-layer CNN, each layer is interpretable as either DNA sequence or TF binding. Thus, the final linear layer directly maps TFs to outputs and can produce a meaningful TF by output matrix with no post-processing. Moreover, the internal parameters of the model, such as trainable pooling or interaction attention, are also directly interpretable and can reveal additional information about mechanisms by which highly parameterized models produce high accuracy predictions. Our approach is inspired by related work that encodes well characterized biochemical systems as neural network functions and optimizes their biochemically interpretable parameters with respect to observed data (Dibaeinia and Sinha, 2021; Liu *et al*., 2020; Tareen and Kinney, 2019). `tiSFM` has several distinct contributions: **(1)**: `tiSFM` is more performant than current state-of-the-art models in the field (Maslova *et al*., 2020) **(2)**: `tiSFM` separates itself from other works by building a “totally” interpretable architecture that is amendable to transparent analysis at each layer, distinct from previous works (Quang and Xie, 2016; Banovich *et al*., 2017; Liu *et al*., 2020) that only rely on initial convolution from PWMs, **(3)**: In our framework, the programmed interpretability in `tiSFM` allows us to offer more extensive homotypic and heterotypic TF-TF interaction insights that are not offered or possible in previous works (Maslova *et al*., 2020) (**Fig. 3**). Overall, our work serves as a general framework that only requires prior knowledge of TF PWMs, the same prior knowledge that would be needed to interpret a normal CNN, post-hoc. In addition, `tiSFM` bridges the gap between high capacity general models and system-specific models with detailed prior specifications, moving deep learning forward with regard to its applications in genomics.

## 2. Methods

### 2.1 Data Processing

We trained `tiSFM` on the ImmGen ATAC-seq dataset (Gal-Oz *et al*., 2019) that consists of assayed open chromatin region (OCR) activity of 81 immune cell-types in the mouse hematopoietic lineage. This dataset was extensively evaluated in a similar deep learning prediction and interpretation task in their paper by Maslova *et al*., 2020. Similarly to this previous work, we focus on predicting correlations across cell-types rather than raw chromatin accessibility. However, rather than changing the form of the loss function as was done in the original study, we achieve a similar effect by z-scoring the activity of each OCR across cell-type and using the standard mean squared error (MSE) loss on the z-scores. Additionally, we focus most of our benchmarking and evaluation on predicting data aggregated over each of the eight main lineage types, rather than the individual cell-types. This is done for two reasons: **(1)** Difficulty — as was demonstrated in the original paper, the within-lineage performance is considerably worse than across-lineages, **(2)** Interpretability evaluation — in many cases, we have substantial knowledge about the TFs that play a role in regulating major lineages, but information about further developmental decisions within a lineage is comparably sparse.

We also train our `tiSFM` on the full dataset; however, rather than using raw data or the z-scores of OCR activities, we focus on OCR activity differences along the cell lineage-tree of hematopoietic differentiation. Specifically, if a cell-type *x* has a parent *P*(*x*), we predict the difference between `OCR_activity`(*x*) and `OCR_activity`(*P*(*x*)). This transformation prevents large differences between lineages from dominating the training objective, instead shifting the focus on developmental transitions. However, this also has the effect of making the problem more difficult and the (*R*^2^) values achieved for all methods are considerably lower than previously reported. We refer to this dataset as “tree-diff”.

The input dataset is summit-centered, and for our model, restricted to the 300 bp centered on the peak summit. We use 862 position weight matrices (PWMs) of mouse TFs from the cisBP database (Weirauch *et al*., 2014a) as weights for our initial convolution kernels.

The cisBP database contains many highly similar PWMs corresponding to same TF or TFs in the same family. While we do not filter for this redundancy in model construction we evaluate it in downstream analyses. We define the similarity of two motifs as *log*_10_ (E-value) where E-values are calculated by TOMTOM (Tanaka *et al*., 2011).

### 2.2 Model training

All models were implemented using PyTorch (Paszke *et al*., 2019). Given the differences in the objectives (see Section 2.1) and train/test splitting, we re-trained the original `AI-TAC` model architecture using our version of the Immgen ATAC-seq dataset, objective, train/validation/test splits, and stopping criteria — allowing for a more direct comparison.

All 512,596 reported genomic regions were divided into 10 approximately similar-sized folds. There are no coordinate overlaps between different peaks and fold assignment was performed at random. In turn, we used 8 folds to form a training set, 1 fold for validation, and 1 fold for testing. We trained models starting with a learning rate of 0.01, decaying by a factor of 0.1 if the validation loss does not improve for 10 epochs; once the learning rate is below 5 × 10^*−*7^ the training stops. We repeated this procedure ten times, with each fold serving as the test set once, and we report the mean performance across test set folds.

`tiSFM` is first trained by freezing the first layer kernel weights to the cisBP PWMs; the kernels are then unfrozen and trained further (for approximately 10 epochs, while monitoring validation improvement as defined above). To encourage sparsity in the final layer, we apply a path algorithm for *L*_1_ or the minimax concave penalty (MCP) regularization via a modified version of the ADAM optimizer; this uses the proximal operators for *L*_1_ and MCP penalties as described in Yun *et al*., 2021.

### 2.3 Model Architecture

All variations of `tiSFM` take one-hot encoded DNA sequence of length *L* as input, which is then propagated through the following network structure:

1. Data transformation.

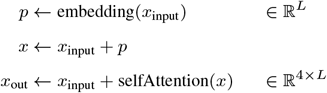

where *p* is broadcast across the rows of *x*_input_ in the second statement, and the output of the self-attention layer matches the dimension of the input. In our models without self-attention we simply skip the last step. These transformations enable the model to account for positional effects (for instance, TFs are known to preferentially bind in the center of ATAC-seq peaks), and to incorporate nucleotide context information.
2. Convolution and scaling. Taking the output from the data transformation block as input convolutions are performed:

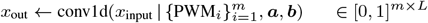

where *m* denotes the number of motifs used, ***a*** ∈ ℝ^*m*^ is a vector of motif-specific scales, and ***b*** ∈ ℝ^*m*^ is a vector of motif-specific offsets. Here and in the following, *L* now represents the length after convolution. This layer computes a convolution of the transformed input with *m* fixed PWM matrices for TF binding sites taken from the CIS-BP database (Weirauch *et al*., 2014b). Although PWMs weights are fixed, every kernel in the convolutional layer (PWM) has two trainable parameters: a scale and a bias, with a sigmoid function as activation. This layer maps raw PWM match scores to [0, 1] in a PWM-specific manner.
3. Trainable pooling. Taking the results from convolution and scaling as input, we learn PWM-specific pooling weights *w*:

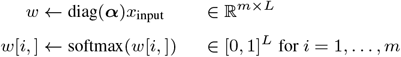

where ***α*** ∈ ℝ^*m*^. We then use these weights to aggregate the input data across the sequence length:

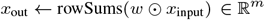

where ⊙ denotes the element-wise product (between learned pooling weights and input sequence). This pooling strategy works similarly to attention pooling, which was popularized by the Enformer model (Avsec *et al*., 2021b). The variable ***α*** acts as a pooling parameter. For ***α***_*i*_ close to zero, pooling weights *w*[*i*,] for PWM_*i*_ will be close to uniform (= average pooling), while for a large ***α***_*i*_ weights will be close to (*m* − 1) zeros and1 one (= max pooling). We note that this trainable pooling strategy, together with the sigmoid activation of the previous step, enables our model to approximate motifmatch counting, which has been shown to be useful in modeling TF-DNA binding (Roider *et al*., 2006). Thus, our model encompasses the entire continuum of possible single motif aggregation strategies ranging from additive binding and occurrence counts under average pooling to saturation effects more accurately modeled by max pooling.
4. Motif interactions. Each motif contribution is multiplied by an interaction-dependent weight:

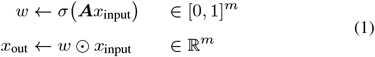

where ***A*** ∈ ℝ^*m×m*^ summarizes motif interactions. This interaction model allows the presence of TF *i* to up- or down-weigh the effective contribution of TF *j*, thus modeling synergistic or saturation interactions across different TFs.
5. OCR activity prediction. With this final, fully connected layer we make a linear prediction of ATAC-seq scores:

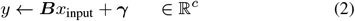

where *y* is the vector of ATAC-seq scores across *c* cell-types, ***B*** ∈ ℝ^*c×m*^ a matrix of regression coefficients, and ***γ*** ∈ ℝ^*c*^ is a vector of intercepts.

### 2.4 Proximal Operators for MCP/ *L*_1_ regularization

To make our model sparse, and therefore more interpretable, we impose sparsity regularization (e.g., *L*_1_, and MCP) on the coefficients of the final layer of our model. Yun *et al*., 2021 formulated the proximal operator for the *L*_1_ regularized ADAM as:

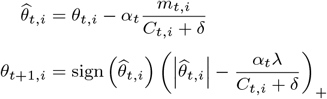

where *C*_*t,i*_ is the preconditioner matrix, *m*_*t,i*_ is the momentum estimate, *α*_*t*_ is the learning rate, *δ* is a small constant added to avoid division by 0, *θ* is the coefficient of the model, and finally *λ* is the hyperparameter for *L*_1_. The operation *f* (*x*) = (*x*)_+_ is defined as *f* (*x*) = max(0, *x*). In a similar vein, they formulated the closed-form solution of projecting through MCP regularization as

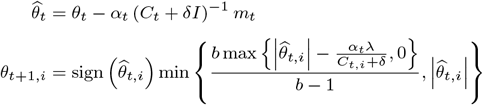

where the parameters are the same with *L*_1_ case, with the addition of *b* which is the second hyperparameter for MCP. While `tiSFM` supports both regularization schemes, results presented in the following use MCP regularization.

## 3. Results

### 3.1 The tiSFM model outperforms a state of the art CNN at OCR activity prediction

An overview of the `tiSFM` architecture can be seen in **Fig. 1A**. Briefly, the `tiSFM` is multi-layer CNN that consists of several elements that have been successfully used across a variety of deep learning models, both across tasks and datasets, such as: convolutional layers, positional embedding, trainable pooling, and self-attention mechanisms. In order to quantify the performance contribution of different parts of the model we implement several variants of this architecture that selectively leave out some of the layers from the full model for comparison (ablation analysis). We also trained all models with fixed kernels (cisBP PWM), and then subsequently relaxed this constraint (fine-tuning).

**Fig. 1.**
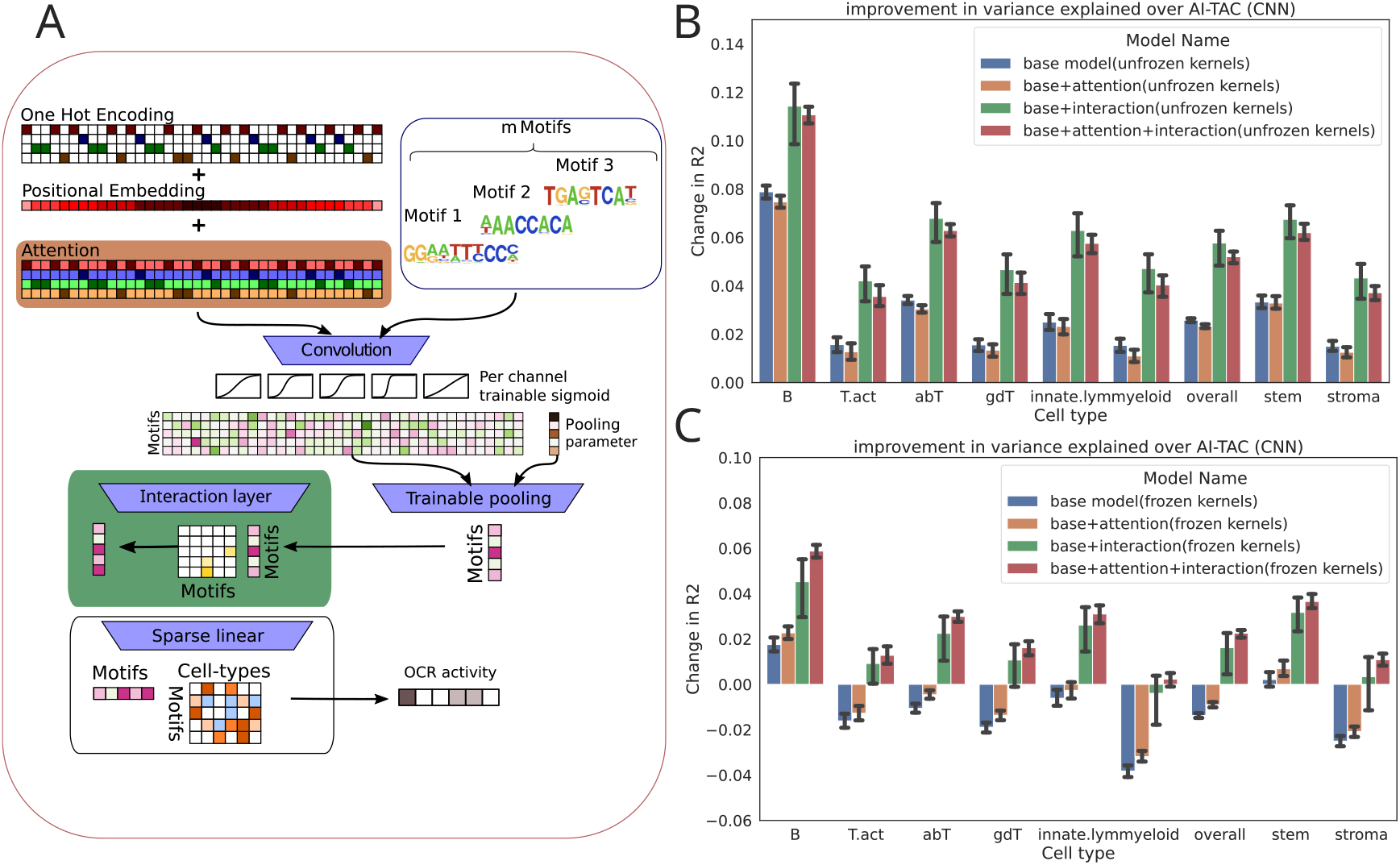
tiSFM improves on prediction accuracy when compared to the current state-of-the-art. (A) A graphical display of the architecture of tiSFM. (B) The improvement of tiSFM, measured via the change in *R*^2^ on the full model; additionally, different iterations of tiSFM were tested with the inclusion/exclusion of certain model components in order to weigh their contribution to the overall performance. (C) similar to (B), but without kernel fine-tuning.

**Figures 1B** and **1C** present the test performance of different model variants evaluated across all 10 cross-validation folds — reported as mean and standard error. Given that we are predicting a continuous activity values, we compare the change in *R*^2^, defined as the difference between `tiSFM`’s and `AI-TAC`’s. Our benchmark, `AI-TAC`, is CNN architecture proposed by Maslova *et al*., 2020, which we retrained according to our the train/validation/test splits and training procedure (see Methods for details). As this architecture was extensively optimized on this dataset, we consider it to be state-of-the-art for this learning task. Notably, our results show that the full `tiSFM` model architecture (represented by red bars in **Figures 1B** and **1C**), with either frozen or unfrozen kernels, outperforms the `AI-TAC` CNN model across all cell-types.

This observation is particularly striking given that `tiSFM` only has 1,037,112 parameters (excluding convolutional kernels, when weights were unfrozen this number drops to 829,752) while `AI-TAC` has nearly three times as many — 2,974,908. This result underscores the importance of incorporating biological prior knowledge in the form of PWMs, and suggests that learning the appropriate kernel weights (or PWMs) is a key source of prediction accuracy. This interpretation is consistent with recent findings demonstrating that a combination of a max-pooled convolution layer and a fully connected linear layer can be competitive with multi-layer CNN networks across a variety of S2F models (Novakovsky *et al*., 2022).

Studying the unfreezing of PWM/kernel weights further supports this line of thought **(Fig. 1C)**, as we find that even the base model, that lacks interaction and attention layers (but still has PWM specific learnable logistic and pooling parameters), can outperform `AI-TAC` after kernel fine-tuning. We note that we also compared these results to a naive linear model, whereby the max-pooled motif scores are fed directly to LASSO regression with the regularization parameter optimized through cross-validation. We find that the base `AI-TAC` model outperforms its linear counterpart demonstrating that the learnable motif calibration and pooling parameters have significant contributions. All model comparisons are detailed in **Table S1**.

While our results strongly indicate that the first layer convolution kernels (which approximate the binding preferences of transcription factors) are the most critical part of our model, other aspects of the model architecture yield additional performance gains. Investigating the impact of attention and/or interaction, we find that both contribute appreciably to model performance, with the biggest contribution coming from modeling TF interactions (**Fig. 1C**). Notably, we observed that this trend holds regardless of whether kernels are frozen or not. Sequence attention, on the other hand, improves performance when kernels are kept fixed and interaction is included; however, performance is somewhat reduced in case kernels are fine-tuned. This suggests that sequence attention may function similarly to kernel fine-tuning by allowing additional flexibility in motif matching. However, we believe fine-tuning PWM weights is the preferable approach, because we find that it does not degrade model interpretability on this data (see below).

### 3.2 Kernel fine-tuning does not alter model interpretability

One concern with the fine-tuning of the kernels is that they will no longer resemble the initial PWM, which would hamper model interpretability. However, in our analysis, we find that this problem does not arise. The PWMs before and after weight unfreezing are very similar, with an average mean squared error (MSE) of 0.04.

Additionally, we examined the pairwise distance between the initial kernels and the final fine-tuned kernels. For 92% of the fine-tuned kernels, the most similar motifs were their pre-fine-tuning counterparts; if a change was detected between the two, the fine-tuned motif often corresponds to a different motif for the same TF. For example this is the case for Dlx1, where the fine-tuned kernel is most similar to the Dlx3 motif. We note that our observation that unfreezing leads to an overall relatively minor change in model parameters (i.e., the difference between fine-tuned and initial kernel weights), but that this change substantially increases performance, further supports the conclusion that getting the first layer kernels “right” is one of the key aspects in building S2F CNNs. However, our analysis also highlights that good kernels can be learned starting from prior knowledge and that fine-tuning is minor and generally does not break interpretability.

### 3.3 tiSFM regression parameters are intrinsically interpretable in terms of lineage specific transcription factors

In the original `AI-TAC` manuscript, the authors employed kernel-PWM matching (also using the mouse CIS-BP database), combined with attribution analysis via DeepLift (Shrikumar *et al*., 2017) to construct a TF-by-cell-type contribution map. In our approach, both of these steps are unnecessary, as the final linear layer already contains this information (parameters ***B*** ∈ R^*c×m*^ in Equation (2)). We visualize the `tiSFM` TF-by-cell-type contribution from a single run of the model in **Fig. 2A**, where the coefficients of the final layer have been normalized to [−1, 1] without breaking sparsity (only non-zero weights are rescaled).

**Fig. 2.**
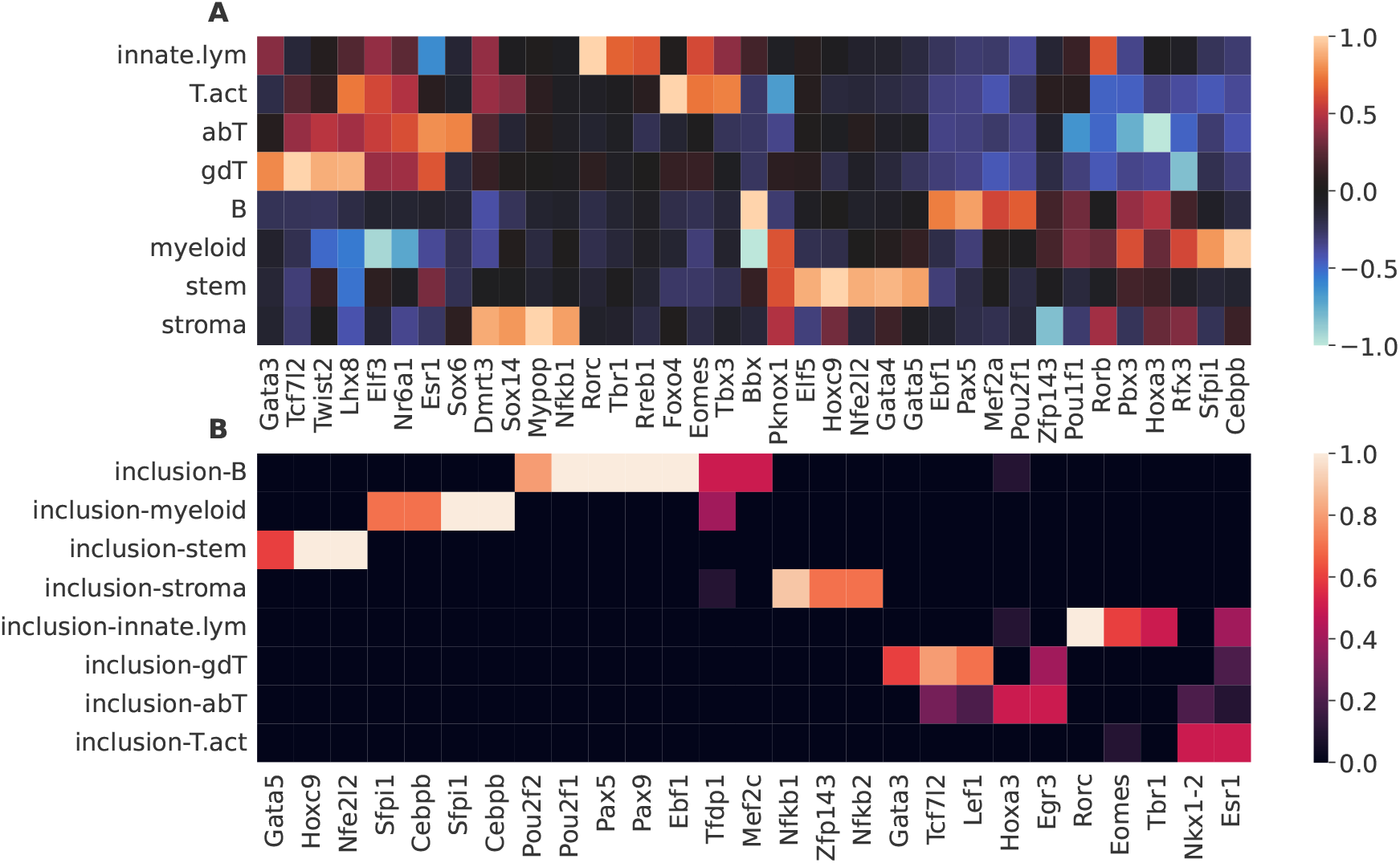
tiSFM consistently finds motifs that are relevant to immune cell differentiation. (A) The final layer of a single fully trained model, after the rows were normalized to [-1, 1], while preserving the sparsity. The 5 motifs with the highest absolute weight for each cell-type are shown. (B) Inclusion ratio was defined as the number of times a motif appeared among the top 10 motifs across the 10 folds in our 10-fold cross-validation procedure, with the highest weights in the final layer for each cell-type. The motifs included in more than 50% of the models are shown here.

**Fig. 3.**
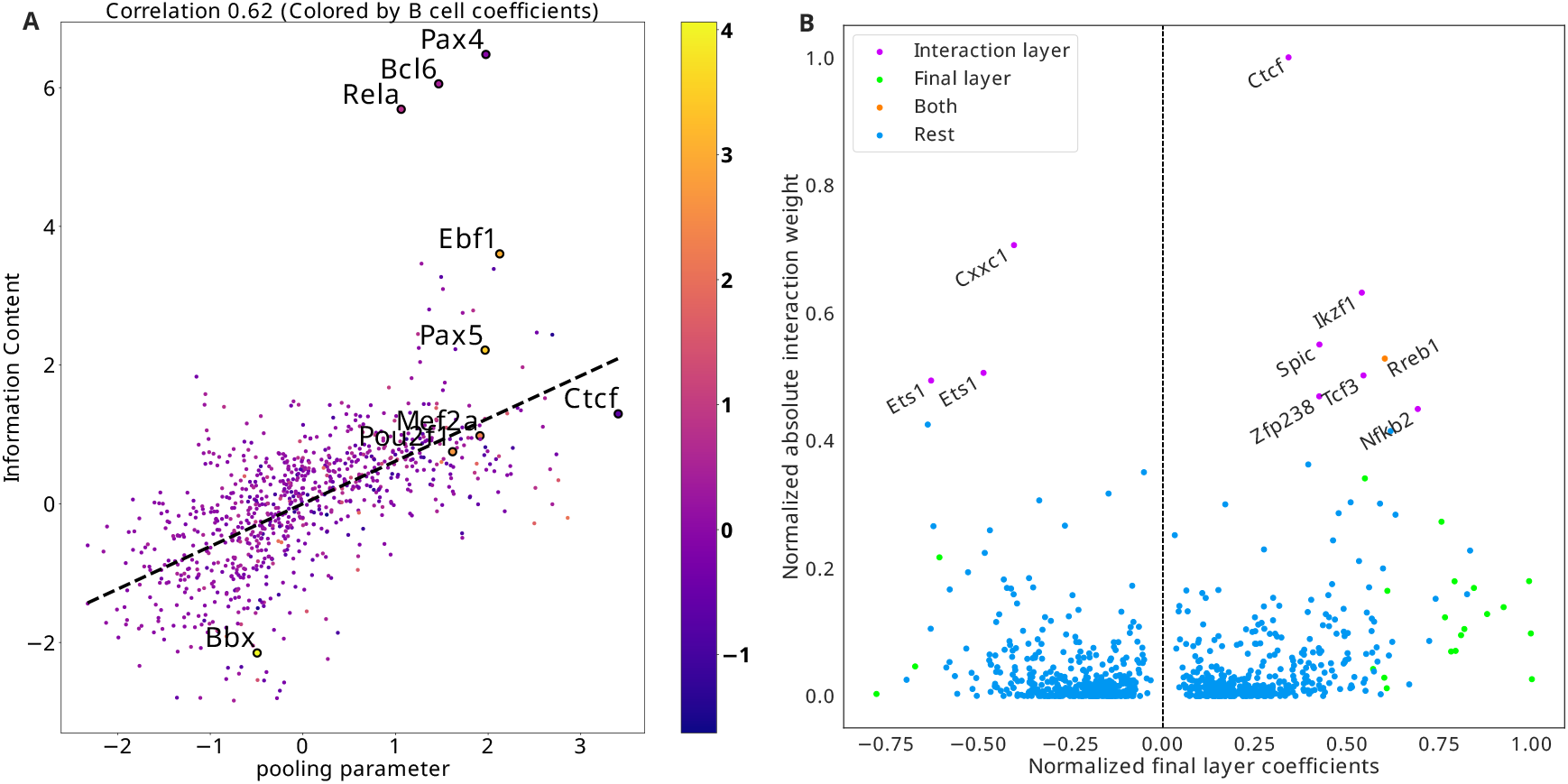
(A) A scatter plot of pooling parameters vs. the information content (IC) calculated from motif PWMs. The color annotation of the points correspond to the final layer coefficients for B-cell, OCR activity prediction. The 5 motifs with the highest absolute weight for B cells are annotated (Ebf1, Pax5, Mef2a, Pou2f1, Bbx) along with some outliers to the pooling/IC trend (Pax4, Bcl6, Rela, Ctcf). (B) A scatter plot contrasting the coefficients in the final linear layer (pooled from absolute values of the cell-type prediction coefficients corresponding to the same TF, then the coefficients are normalized to [*−*1, 1], range preserving the sparsity) with total contribution to the interaction layer. The two are notably different, only one TF influences the other TFs while contributing to the cell-type prediction significantly, as some TFs contribute far more to the interaction matrix but have near 0 linear coefficients.

We find that `tiSFM` effectively highlights known regulators of immune cell differentiation. For B cells the model selects Pax5 (with Pax9, an additional TF in the family) and Ebf1, which are known as *the* key early regulators of B-cell lineage commitment (Somasundaram *et al*., 2021; Hagman *et al*., 2011). The model also selects Irf4 which is again consistent with known roles of Irf4/Irf8 in B cell development (Shukla and Lu, 2014).

`tiSFM` also highlights Cebpb and Sfpi1 (a.k.a. Spi1/PU.1) for myeloid cells and Rorc and Eomes for innate lymphoid cells. These TFs are likewise well-characterized master regulators of their respective lineages (Suñer *et al*., 2022; Hoorweg *et al*., 2012; Kiekens *et al*., 2021). Importantly, these TFs were also highlighted in the original `AI-TAC` analysis, and overall the `AI-TAC` and `tiSFM` TF by cell-type matrices are highly similar; an important difference, though, is that for the `tiSFM` model these are simply model parameters and no additional post-hoc calculations were required.

To assess the consistency of our approach, we used the cross-validation folds from model training to calcuate a per-motif inclusion ratio — defined as the number of times a motif appeared in the top 10 motifs with the highest absolute weights from each cell-type. Several of the motifs reported consistently appeared in the top 10. Indeed, all the well known lineage regulators discussed above (Pax5, Ebf1, Cebp, Sfpi1, and Rorc) have an inclusion ratio of 1 (**Fig 2B**), suggesting that TFs with similarly high inclusion ratio are likely true positives. One such TF is Hoxc9, which is highlighted by `tiSFM` as being important for hematopoietic stem cells (HSCs). While no such function for Hoxc9 has been established, Hoxa9 is a highly similar motif (cisBP database the Hoxa9 consensus is a subset of the Hoxc9 consensus) that is known to play a critical role in hematopoiesis. Defects in Hoxa9 lead to an inability of HSCs to repopulate an irradiated host (Lawrence *et al*., 2005). On the other hand, over-expression of Hoxa9 increases the efficiency of hematopoiesis (Ramos-Mejía *et al*., 2014).

We also note that the consistency of top transcription factors for T cells is notably lower (**Fig 2B**); nevertheless, among the ones shown, we find several that are canonically associated with maintaining T cell function, including Lef1, Tcf7l2 (Tcf7 family), Egr3, and Gata3 (Xing *et al*., 2016; Shan *et al*., 2021; Ho *et al*., 2009; Li *et al*., 2012). Notably, while Gata3 has a known role in early T cell development, the rest of these TFs are associated with maintaining the function of specific T cell sub-types, not necessarily with establishing T cell identity. This result mirrors the original `AI-TAC` finding and conclusion that the observed weak T-cell attribution implied that the T-cell lineage was a fall back plan that did not require specific regulators. Rothenberg, 2011 supports this notion with experimental evidence. TF coefficients and consistency for alternative versions of `tiSFM` is shown in **Figures S1** and **S2**.

### 3.4 Internal tiSFM parameters have intuitive biological interpretations

We have demonstrated that the final linear layer of our `tiSFM` model is interpretable and highly consistent with known biology. However, other internal parameters of the model also have biological and biochemical interpretations.

For example, our model includes a trainable pooling parameter that can interpolate between max and average pooling (see Methods). As the input to the pooling is passed through a sigmoid activation with TF specific scaling and offset, the trainable pooling step can potentially count binding events, rather than simply retaining the maximal motif match. From a biochemical perspective we expect that TFs with weak protein DNA interactions to be more dependent on multiple instances of a binding motif. As such, we expect that motifs with relatively lower information content (as proxy for low-sequence specificity) would prefer average pooling. This is indeed what we observe by plotting motif information content against pooling parameters in **Fig. 3A**. There we see that the information content and pooling parameter are strongly correlated. We note that Ctcf has the largest pooling parameter, indicating that the strength of the strongest Ctcf site is the most useful featurization of local Ctcf activity. In **Fig. 3A**, weights of the final model layer for B-cells are color coded; no trend is observed, indicating that the association of motif information content and pooling parameter is independent of final layer weights. We highlighted 5 motifs with the highest absolute weight for B cell prediction: Ebf1, Pax5, Mef2a, Pou2f1, Bbx; this was done along with motifs that were outliers relative to the overall trend between information content and pooling parameter (Rela, Bcl6, Pax4).

**Fig. 3B** displays the total interaction influence of motifs, computed as the column sums of the absolute values of the matrix ***A*** ∈ R^*m×m*^ of the interaction layer (Equation (1)). This layer enables the final pooled score of one motif to influence the score of another motif, while maintaining directionality (i.e., ***A*** is not necessarily symmetric). Plotting interaction influence against the total final layer influence (defined as the maximum of the absolute value of coefficients across cell-types), we find that the top interaction TFs are not the same as the ones with the most influence on the final output. Indeed, Ctcf has the largest overall interaction influence, but does not appear as a top TF for any cell-type. This is consistent with known biology as Ctcf is an important regulator of 3D genome organization and is likely to be expressed in every cell-type and thus should not be predictive of cell-type identity.

The other two motifs that score highly for interaction, Cxxc1 and Ikzf1, also have low linear layer coefficients. While these TFs do have roles in specifying cell-fate, they also have short and thus low-affinity recognition sequences and partake in complex interaction networks with other TFs that increase their binding affinity (Kiuchi *et al*., 2021; Marke *et al*., 2018).

### 3.5 Sparsity constraints improve performance for most cell-types and increase interpretability of tiSFM

To induce sparsity in `tiSFM`’s regression coefficients in the final layer, we make use of the path algorithm first put forward by Park and Hastie, 2007 in their seminal work. Specifically, we utilized the proximal operators for *L*_1_ and MCP to enforce sparsity on the coefficient matrix ***B*** in Equation (2) to select a few motifs that are crucial for prediction of cell-type-specific OCR activity. The path-finding algorithm starts with the fully trained model and progressively increases the intensity of the penalty (by increasing MCP’s hyperparameter, *λ*, while keeping the second hyperparameter fixed at *b* = 3), starting at each step from the model that was converged on at the previous step. Results are summarized in **Fig. 4**.

**Fig. 4.**
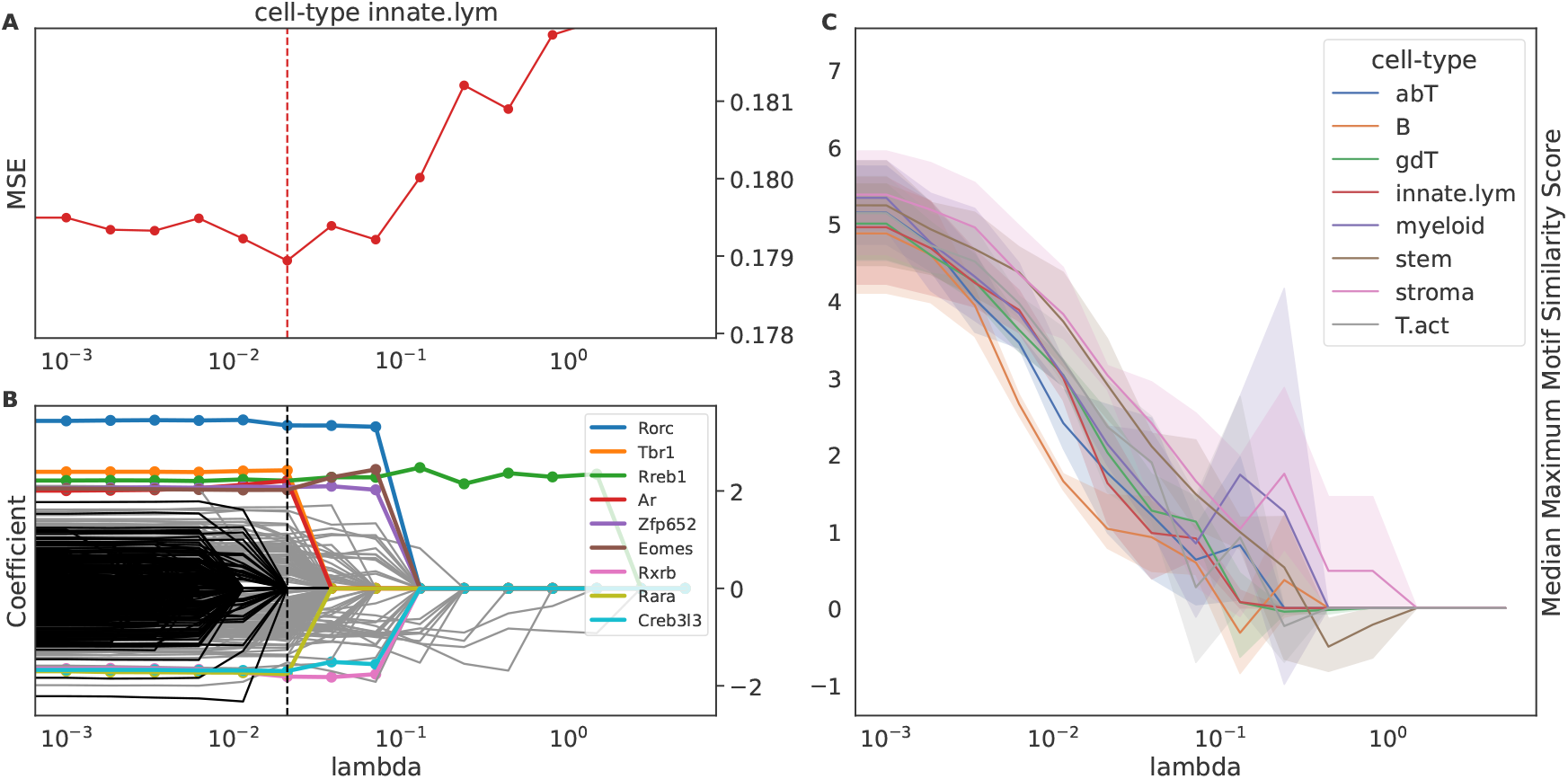
Sparsity constraints improve the performance for the most cell-types and increase interpretability of the tiSFM. (A) MSE vs the MCP hyperparameter (the second hyperparameter is fixed at 3). The lines for every cell-type and the overall performance are assigned a color. The vertical line indicates the hyperparameter that resulted in the best performance for the corresponding cell-type. (B) Coefficients vs the MCP hyperparameter path plot. The best model in terms of MSE is marked by a dashed vertical line. Colored, are the top 9 motifs with the highest absolute coefficients from the best model, and the black lines are the motifs with 0 coefficient in the best model. (C) Median motif redundancy as a function of regularization. We observe that as expected increasing regularization decreases motif redundancy (see Methods for motif similarity calculation). Path plots for other cell-types are in Figure S3.

While we observe that the sparsity does not substantially improve model performance (**Fig. 4A**), by enforcing it we can alleviate motif redundancy present among PWMs and achieve sparser, more interpretable models that can perform at least as well as the full model. Sparser representations allow us to focus on consistently reappearing, distinct motifs, indicating their importance in contributing to the `tiSFM` model. **Fig. 4B** shows the regularization path for innate lymphocytes, similar plots for other lineages are in supplemental **Figure S3**. We observe that choosing the best-performing model induces sparsity (many PWMs’ contribution is zero); PWMs with the largest coefficients are colored, and we note that Rorc is consistently selected by the path algorithm as one of the proteins that contributes most to the epigenetic characteristics of the innate lymphoid cells.

We also evaluate how regularization impacts the inclusion of redundant motifs. Specifically, for each TF with a non-zero coefficient in the final layer of the model we compute the maximum TOMTOM similarity score (see Methods) to all other included motifs. We observe that increasing the sparsity results in the selection of more distinct transcription factors, as shown by the decreasing trends in the median maximum similarity score, as illustrated in **Fig. 4C**.

### 3.6 tiSFM identifies key regulatory events along differentiation trajectories

For our analyses, we use `tiSFM` to analyze OCR activity changes between parents and children in the hematopoietic lineage-tree, which we term “tree-diff” (see Methods). This problem is considerably more challenging than predicting open chromatin in cell-types per se. For instance, using lineage-aggregated data (i.e., pooling all cell-types within a lineage), all models are able to achieve a mean (across cell-types) *R*^2^ of about 0.2 or a Pearson correlation of about 0.4 with `AI-TAC`’s *R*^2^ range being 0.1 (stem cell) to 0.31 (myeloid). On the “tree-diff” data the corresponding values are around 0.05 (*R*^2^) and 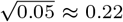 (correlation) (see Table S2 for all *R*^2^ values). Moreover, there are large differences in performance, with some changes having *R*^2^ values near 0. We found that the predictability, as measured by *R*^2^ was strongly related to the amount of variance along the transitions edge (Figure S4). Generally, developmental transitions that induce only a small amount of epigenetic change (low variance) are harder to predict.

We focus our analysis on those lineage differences that had *R*^2^ *>* 0.05 (equivalently, correlation *>* 0.22). For visualization in **Fig. 5** these edges are scaled according to *R*^2^ and color-coded according to the transition identity. The same colors are used to indicate TF importance (as inferred from final model layer) for each well-predicted transition (**Fig. 5 B**). The TF importance heatmap is annotated with the color of the corresponding tree edge.

**Fig. 5.**
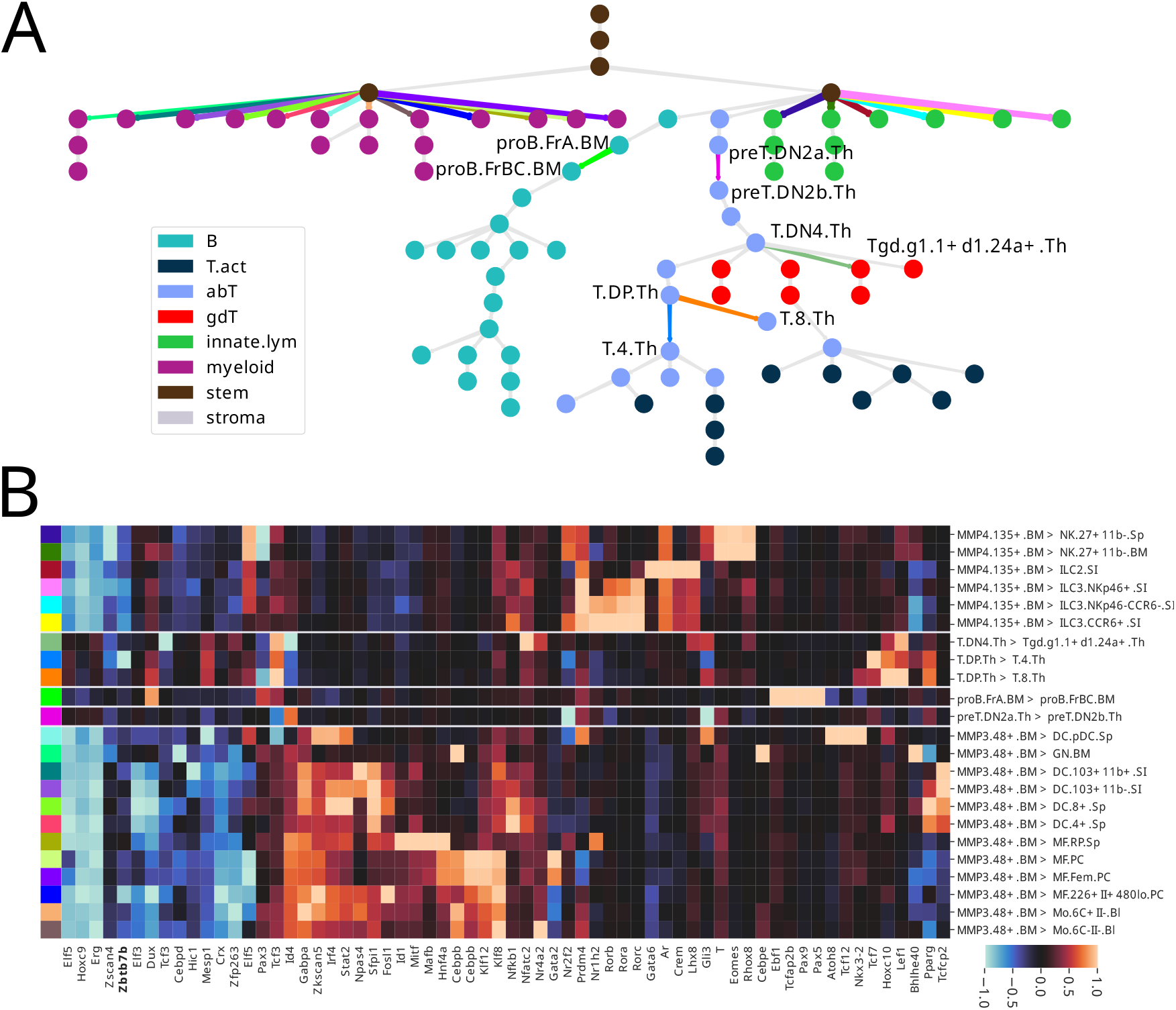
tiSFM predictions and TF contributions applied to the problem of predicting differentiation transitions. (A) The model predicts the output corresponding to each edge along the differentiation tree computed as the difference between child and parent. Edges are scaled according to *R*^2^ and those with a value of *>* 0.05 are selected for TF contribution analysis depicted in the color matched heatmap (B). Highly predictable transitions in the lymphocyte lineage are highlighted on the lineage tree with cell-type identifiers. The heatmap includes motifs that are among top 5 with the highest absolute coefficients for at least one target.

Despite relatively few edges with high quality predictions, we find striking consistency with known biology among the transitions that we predict well. The lineage transitions that stand out and are modeled most clearly are those leading to NK, ILC, and myeloid subtypes from their respective common progenitors. The TF influence map for these transitions are highly correlated — this indicates that we capture largely general development, and only to a lesser extent, differentiation into specific sub-types. For the NK, ILC, and myeloid transitions, TF contributions mirror closely the ones we observed when analyzing lineage aggregated data.

There is comparatively few well predicted transitions in the lymphocyte portion of the tree. Strikingly though, one of the transitions that is predicted well is the CD4/CD8 split. In fact, for this transition, `tiSFM` highlights Zbtb7b as a key regulator of CD4 cells — a result not observed when the T cell lineage is predicted in aggregate. Zbtb7b (often referred to in the literature as ThPOK) is a repressor that is well known to be essential for CD4^+^ T cell differentiation (Basu *et al*., 2021). As Ztb7b acts by repressing the alternative CD8 lineage, the observed negative coefficient is consistent with known biology. Our “tree-diff” analysis also recovers Pax5 and Ebf1 as known master B-cell regulators; interestingly, though, it places their influence at a specific developmental transition denoted as “proB.FrBC.BM-proB.FrA.BM” thus pinpointing the time point at which these regulators induce lineage commitment.

## 4 Discussion

Non-coding regulatory grammar, encoded in the nucleotide sequence of genomes, mediates genome function through complex interactions between transcription factors and their DNA binding motifs. Here we propose `tiSFM`, a novel CNN-based architecture for predicting functional genomic readouts directly from sequence. Our approach is capable of matching and exceeding the performance of state-of-the-art architectures, while at the same time providing immediately interpretable model parameters.

Using a dataset of hematopoietic development we show that our model re-discovers essential regulators previously highlighted by a much more complex post-hoc interpretation approach (Maslova *et al*., 2020), without the need for any additional analysis. We also demonstrate that other internal model parameters, such as the trainable pooling parameters and TF-TF interactions are interpretable in terms of biochemical principles and known biology. Furthermore, applying our model to above-mentioned data, and considering accessibility changes across the hematopoietic lineage-tree, we identify specific developmental transitions that are highly predictable from sequence; this indicates that they correspond to large chromatin remodeling events that are driven by local TF activity.

`tiSFM` has fewer parameters compared with current state-of-the-art models like `AI-TAC`; instead, it relies on prior knowledge regarding sequence preferences of TFs. A possible shortcoming of our approach is that it is somewhat limited to sequence patterns present in the motif database we use. Therefore, it could miss important signals from currently unknown motifs, a limitation that is not shared by other methods that learn convolution weights from scratch. However, since `tiSFM` outperforms the state-of-the-art from-scratch model architecture on this dataset, using prior motif knowledge does not appear to be a limiting factor, performance-wise. Indeed, the observation that frozen kernels can outperform trainable kernels highlights that learning the right TF binding preferences is critical to good performance in sequence-to-function modeling. This result further suggests that subsequent CNN layers in deep CNNs mostly perform kernel refinement rather than compute complex regulatory grammar.

Overall, our work demonstrates progress towards a model that has high predictive capacity but is also interpretable in the context of current knowledge. However, significant challenges remain. First, it is not always possible to identify a true regulator from the PWM relevance, because many TFs come in families with highly similar PWMs. For example, our model highlighted Hoxc9 as important for hematopoietic stem cells, while the correct protein involved is most likely Hoxa9. This problem is not specific to our architecture and could be resolved (at least partially) by cross-referencing putative regulators with genes that show RNA or protein expression in the cell-type or sample of interest. Second, we show that explicitly accounting for TF-TF interactions increases model performance, and that the interaction coefficients we recover are consistent with known biology; nevertheless, the interaction model could be made more biochemically relevant. For example, our current interaction layer is applied after global pooling and thus does not consider distance or orientation between TF motifs in the input sequence. A natural extension would be to apply self-attention before the global pooling layer; however, we found that this approach significantly degraded final layer interpretability as measured by occurrence of known lineage

TFs and fold-reproducibility for the data we studied. The interaction layer presented here is also not cell-type specific, which is an important limitation, because biochemical interactions can only occur if both species are present, but not all TFs are expressed in all cell-types. Expanding on the TF-TF interaction model to take into account cell-type specificity and positional effects will be the subject of future efforts. Nevertheless, taken everything together `tiSFM` presents a step forward in interpretable sequence-to-function modeling and can readily be applied to contexts other than modeling open chromatin during blood cell differentiation.

## Acknowledgments

We acknowledge Sara Mostafavi for help with the Immgen data and helpful technical discussions.

## References

Alipanahi, B. et al. (2015). Predicting the sequence specificities of DNA- and RNA-binding proteins by deep learning. Nat. Biotechnol., 33, 831– 838.

Avsec, Ž. et al. (2021a). Base-resolution models of transcription-factor binding reveal soft motif syntax. Nature Genetics, 53, 354–366.

Avsec, et al. (2021b). Effective gene expression prediction from sequence by integrating long-range interactions. Nature Methods, 18(10), 1196– 1203. Number: 10 Publisher: Nature Publishing Group.

Banovich, N. E. et al. (2017). Impact of regulatory variation across human iPSCs and differentiated cells. Genome Research.

Basu, J. et al. (2021). Essential role of a ThPOK autoregulatory loop in the maintenance of mature CD4+ T cell identity and function. Nat. Immunol., 22, 969–982.

Dibaeinia, P. and Sinha, S. (2021). Deciphering enhancer sequence using thermodynamics-based models and convolutional neural networks. Nucleic Acids Research, 49(18), 10309–10327.

ENCODE Project Consortium (2012). An integrated encyclopedia of DNA elements in the human genome. Nature, 489(7414), 57–74.

Gal-Oz, S. T. et al. (2019). ImmGen report: sexual dimorphism in the immune system transcriptome. Nature Communications, 10(1), 4295.

Hagman, J. et al. (2011). B lymphocyte lineage specification, commitment and epigenetic control of transcription by early b cell factor 1. In Current Topics in Microbiology and Immunology, pages 17–38. Springer Berlin Heidelberg.

Ho, I.-C. et al. (2009). GATA3 and the T-cell lineage: essential functions before and after T-helper-2-cell differentiation. Nat. Rev. Immunol., 9(2), 125.

Hoorweg, K. et al. (2012). Functional differences between human NKp44- and NKp44 RORC innate lymphoid cells. Frontiers in Immunology, 3.

Kelley, D. R. et al. (2016). Basset: learning the regulatory code of the accessible genome with deep convolutional neural networks. Genome Res., 26(7), 990–999.

Kiekens, L. et al. (2021). T-BET and EOMES accelerate and enhance functional differentiation of human natural killer cells. Frontiers in Immunology, 12.

Kiuchi, M. et al. (2021). The Cxxc1 subunit of the Trithorax complex directs epigenetic licensing of CD4+ T cell differentiation. J. Exp. Med., 218(4). 10 Balci et al.

Koo, P. K. and Eddy, S. R. (2019). Representation learning of genomic sequence motifs with convolutional neural networks. PLoS Comput. Biol., 15(12), e1007560.

Lawrence, H. J. et al. (2005). Loss of expression of the hoxa-9 homeobox gene impairs the proliferation and repopulating ability of hematopoietic stem cells. Blood, 106(12), 3988–3994.

Li, S. et al. (2012). The Transcription Factors Egr2 and Egr3 Are Essential for the Control of Inflammation and Antigen-Induced Proliferation of B and T Cells. Immunity, 37(4), 685–696.

Liu, Y. et al. (2020). Fully interpretable deep learning model of transcriptional control. Bioinformatics, 36(Supplement_1_), 499 − −i507.

Marke, R. et al. (2018). The many faces of IKZF1 in B-cell precursor acute lymphoblastic leukemia. Haematologica, 103(4), 565.

Maslova, A. et al. (2020). Deep learning of immune cell differentiation. Proc. Natl. Acad. Sci. U.S.A., 117(41), 25655–25666.

Novakovsky, G. et al. (2022). ExplaiNN: interpretable and transparent neural networks for genomics. bioRxiv, page 2022.05.20.492818.

Park, M. Y. and Hastie, T. (2007). L1-regularization path algorithm for generalized linear models. Journal of the Royal Statistical Society: Series B (Statistical Methodology), 69(4), 659– 677._eprint: https://onlinelibrary.wiley.com/doi/pdf/10.1111/j.1467-9868.2007.00607.x.

Paszke, A. et al. (2019). Pytorch: An imperative style, high-performance deep learning library.

Quang, D. and Xie, X. (2016). DanQ: a hybrid convolutional and recurrent deep neural network for quantifying the function of DNA sequences. Nucleic Acids Res., 44(11), e107.

Ramos-Mejía, V. et al. (2014). HOXA9 promotes hematopoietic commitment of human embryonic stem cells. Blood, 124(20), 3065–3075.

Roider, H. G. et al. (2006). Predicting transcription factor affinities to DNA from a biophysical model. Bioinformatics, 23(2), 134–141.

Rothenberg, E. V. (2011). T cell lineage commitment: identity and renunciation. Journal of immunology (Baltimore, Md. : 1950), 186(12), 6649.

Shan, Q. et al. (2021). Tcf1 and Lef1 provide constant supervision to mature CD8+ T cell identity and function by organizing genomic architecture. Nat. Commun., 12(5863), 1–20.

Shrikumar, A. et al. (2017). Learning important features through propagating activation differences. In International conference on machine learning, pages 3145–3153. PMLR.

Shukla, V. and Lu, R. (2014). IRF4 and IRF8: Governing the virtues of B Lymphocytes. Frontiers in biology, 9(4), 269.

Somasundaram, R. et al. (2021). EBF1 and PAX5 control pro-b cell expansion via opposing regulation of the imyc/i gene. Blood, 137(22), 3037–3049.

Suñer, C. et al. (2022). Macrophage inflammation resolution requires CPEB4-directed offsetting of mRNA degradation. eLife, 11.

Tanaka, E. et al. (2011). Improved similarity scores for comparing motifs. Bioinformatics, 27(12), 1603.

Tareen, A. and Kinney, J. B. (2019). Biophysical models of cis-regulation as interpretable neural networks. arXiv.

Weirauch, M. T. et al. (2014a). Determination and inference of eukaryotic transcription factor sequence specificity. Cell, 158(6), 1431–1443.

Weirauch, M. T. et al. (2014b). Determination and inference of eukaryotic transcription factor sequence specificity. Cell, 158(6), 1431–1443.

Xing, S. et al. (2016). Tcf1 and Lef1 transcription factors establish CD8+ T cell identity through intrinsic HDAC activity. Nat. Immunol., 17(6), 695.

Yun, J. et al./person-group>. (2021). Adaptive Proximal Gradient Methods for Structured Neural Networks. In M. Ranzato, A. Beygelzimer, Y. Dauphin, P. S. Liang, and J. W. Vaughan, editors, Advances in Neural Information Processing Systems, volume 34, pages 24365–24378. Curran Associates, Inc.

Zhou, J. and Troyanskaya, O. G. (2015). Predicting effects of noncoding variants with deep learning-based sequence model. Nat. Methods, 12(10), 931–934.

