## Supplemental Material for "An intrinsically interpretable neural network architecture for sequence to function learning"

| | abT | B | gdT | innate.lym | myeloid | stem | stroma | T.act |
|---|---|---|---|---|---|---|---|---|
| model names | | | | | | | | |
| base model(frozen kernels) | $0.196 \pm 0.003$ | $0.149 \pm 0.005$ | $0.145 \pm 0.004$ | $0.163 \pm 0.006$ | $0.273 \pm 0.004$ | $0.108 \pm 0.005$ | $0.235 \pm 0.004$ | $0.142 \pm 0.005$ |
| base model(unfrozen kernels) | $0.240 \pm 0.003$ | $0.210 \pm 0.005$ | $0.180 \pm 0.004$ | $0.195 \pm 0.006$ | $0.326 \pm 0.004$ | $0.139 \pm 0.005$ | $0.275 \pm 0.004$ | $0.174 \pm 0.005$ |
| base+attention(frozen kernels) | $0.202 \pm 0.003$ | $0.154 \pm 0.005$ | $0.151 \pm 0.004$ | $0.167 \pm 0.006$ | $0.279 \pm 0.004$ | $0.113 \pm 0.006$ | $0.239 \pm 0.004$ | $0.146 \pm 0.005$ |
| base+attention(unfrozen kernels) | $0.237 \pm 0.003$ | $0.206 \pm 0.004$ | $0.178 \pm 0.004$ | $0.193 \pm 0.006$ | $0.322 \pm 0.004$ | $0.139 \pm 0.004$ | $0.272 \pm 0.003$ | $0.171 \pm 0.006$ |
| base+attention+interaction(frozen kernels) | $0.236 \pm 0.004$ | $0.190 \pm 0.005$ | $0.180 \pm 0.005$ | $0.201 \pm 0.007$ | $0.313 \pm 0.005$ | $0.142 \pm 0.006$ | $0.270 \pm 0.005$ | $0.171 \pm 0.006$ |
| base+attention+interaction(unfrozen kernels) | $0.269 \pm 0.004$ | $0.242 \pm 0.006$ | $0.206 \pm 0.007$ | $0.227 \pm 0.007$ | $0.351 \pm 0.007$ | $0.168 \pm 0.006$ | $0.297 \pm 0.005$ | $0.194 \pm 0.008$ |
| base+interaction(frozen kernels) | $0.229 \pm 0.018$ | $0.177 \pm 0.024$ | $0.175 \pm 0.018$ | $0.196 \pm 0.018$ | $0.307 \pm 0.021$ | $0.138 \pm 0.013$ | $0.263 \pm 0.023$ | $0.168 \pm 0.014$ |
| base+interaction(unfrozen kernels) | $\mathbf{0.274 \pm 0.015}$ | $\mathbf{0.246 \pm 0.024}$ | $\mathbf{0.211 \pm 0.015}$ | $\mathbf{0.232 \pm 0.016}$ | $\mathbf{0.358 \pm 0.015}$ | $\mathbf{0.173 \pm 0.012}$ | $\mathbf{0.303 \pm 0.013}$ | $\mathbf{0.201 \pm 0.013}$ |
| AI-TAC | $0.208 \pm 0.006$ | $0.135 \pm 0.008$ | $0.165 \pm 0.009$ | $0.168 \pm 0.008$ | $0.313 \pm 0.006$ | $0.103 \pm 0.006$ | $0.259 \pm 0.006$ | $0.160 \pm 0.009$ |
| Lasso Regression | $0.120 \pm 0.008$ | $0.091 \pm 0.007$ | $0.087 \pm 0.008$ | $0.102 \pm 0.006$ | $0.175 \pm 0.005$ | $0.061 \pm 0.006$ | $0.162 \pm 0.008$ | $0.087 \pm 0.006$ |

Table S1: We computed the $R^2$ values for each model, and all models underwent training and testing using the identical 10 data folds.

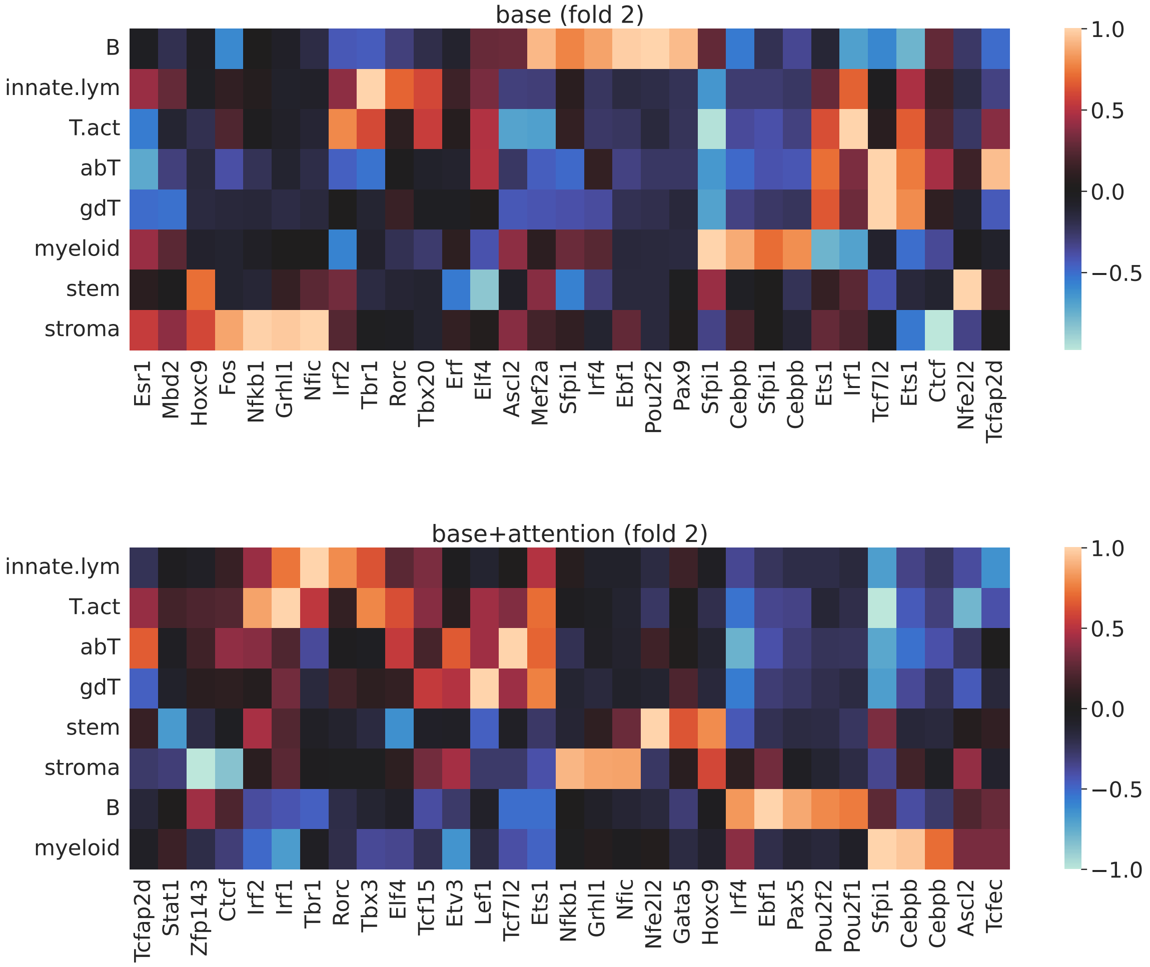


Figure S1: The final-layer coefficients captured by different versions of the tiSFM model.

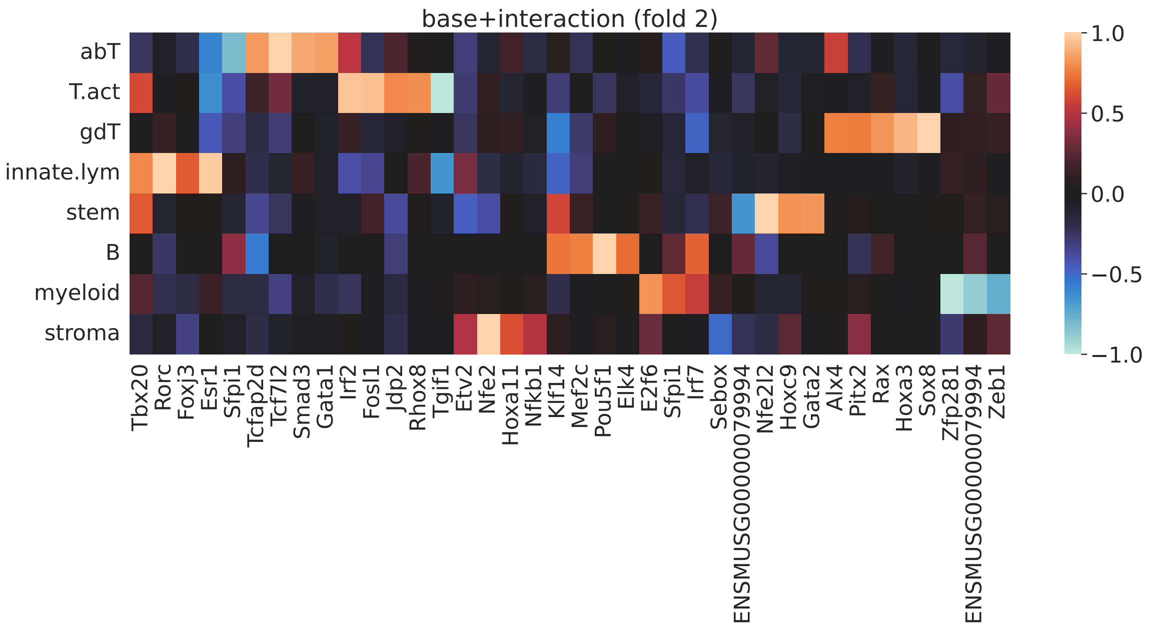


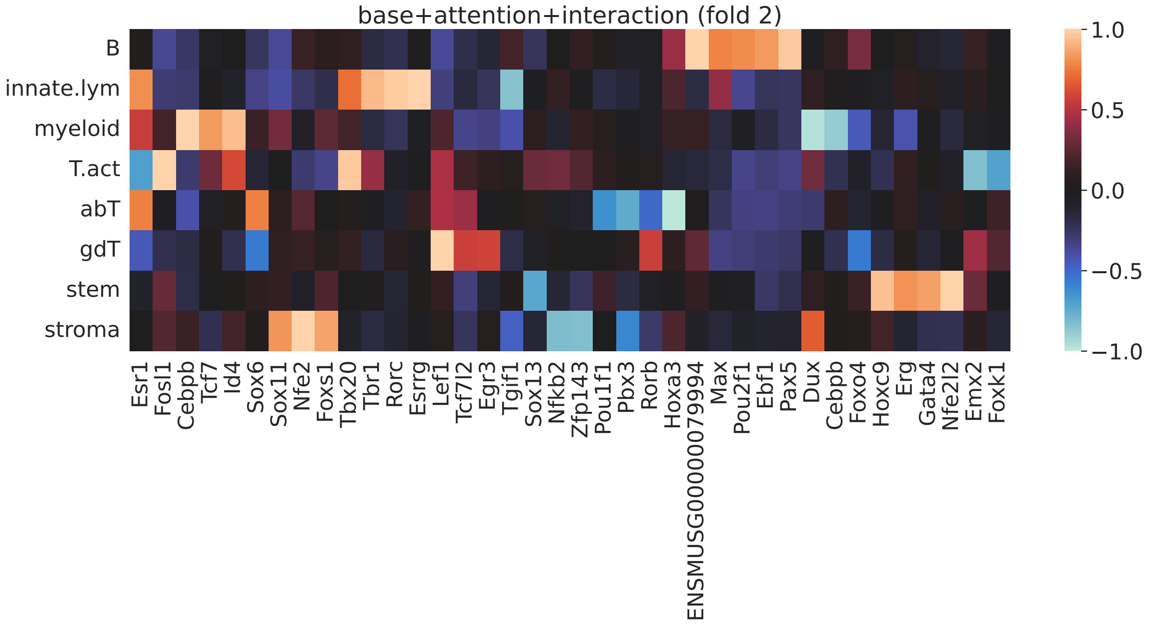


Figure S1: The final-layer coefficients captured by different versions of the tiSFM model (cont.).

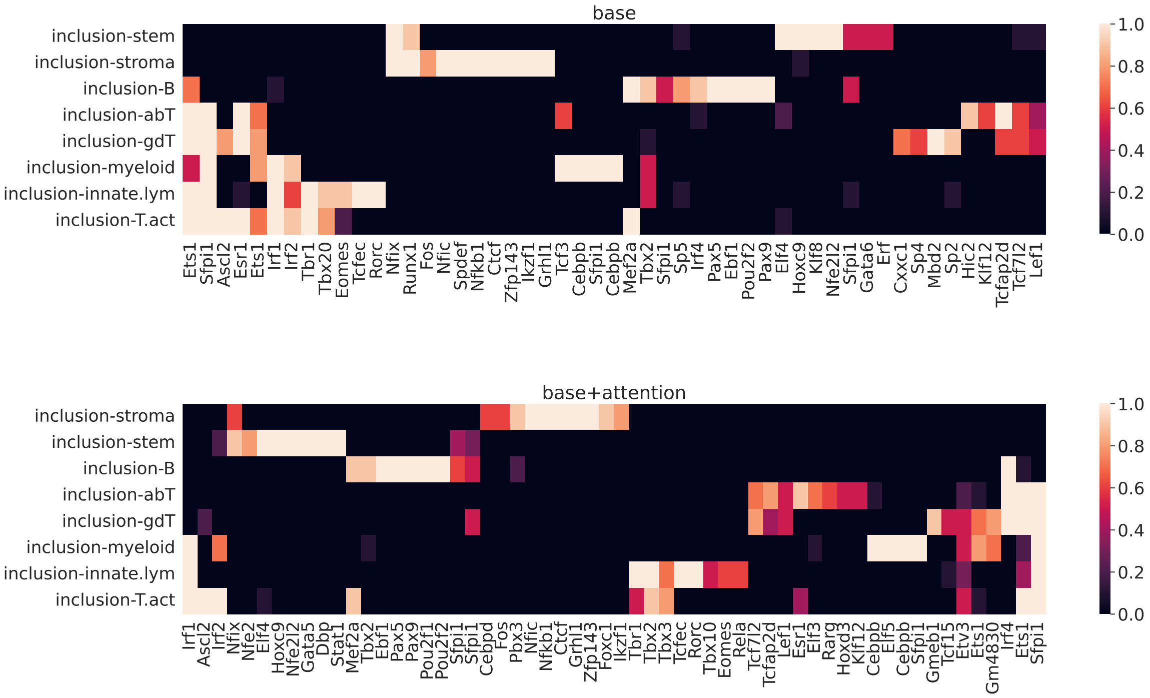


Figure S2: Heatmaps dictating inclusion across the total amount of folds tested.

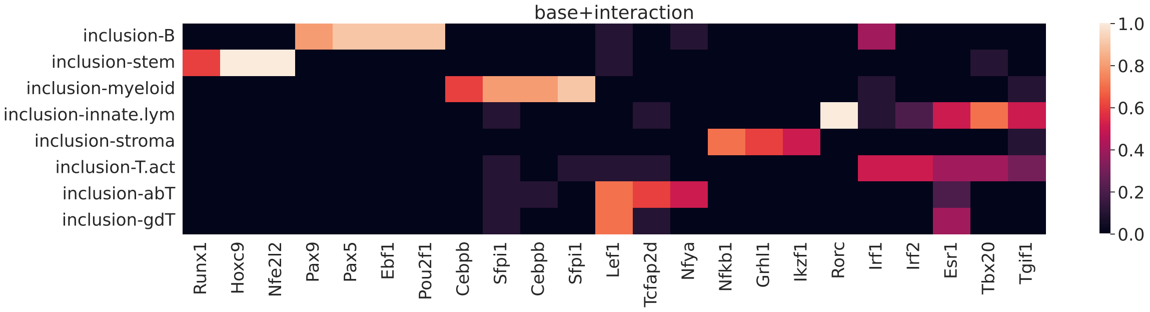


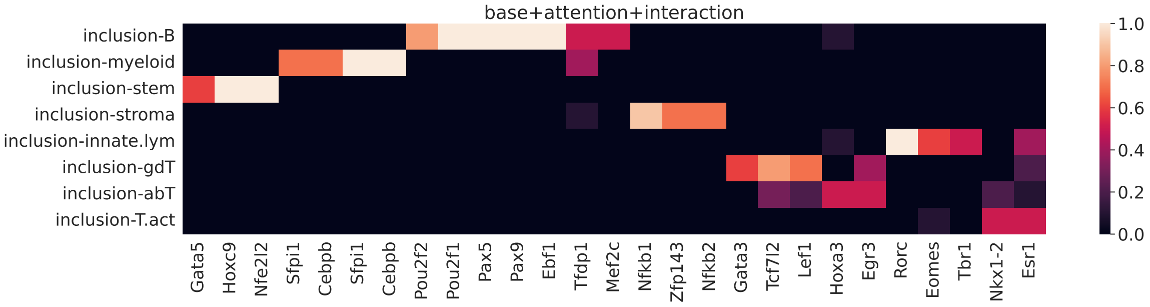


Figure S2: Heatmaps dictating inclusion across the total amount of folds tested (cont.).

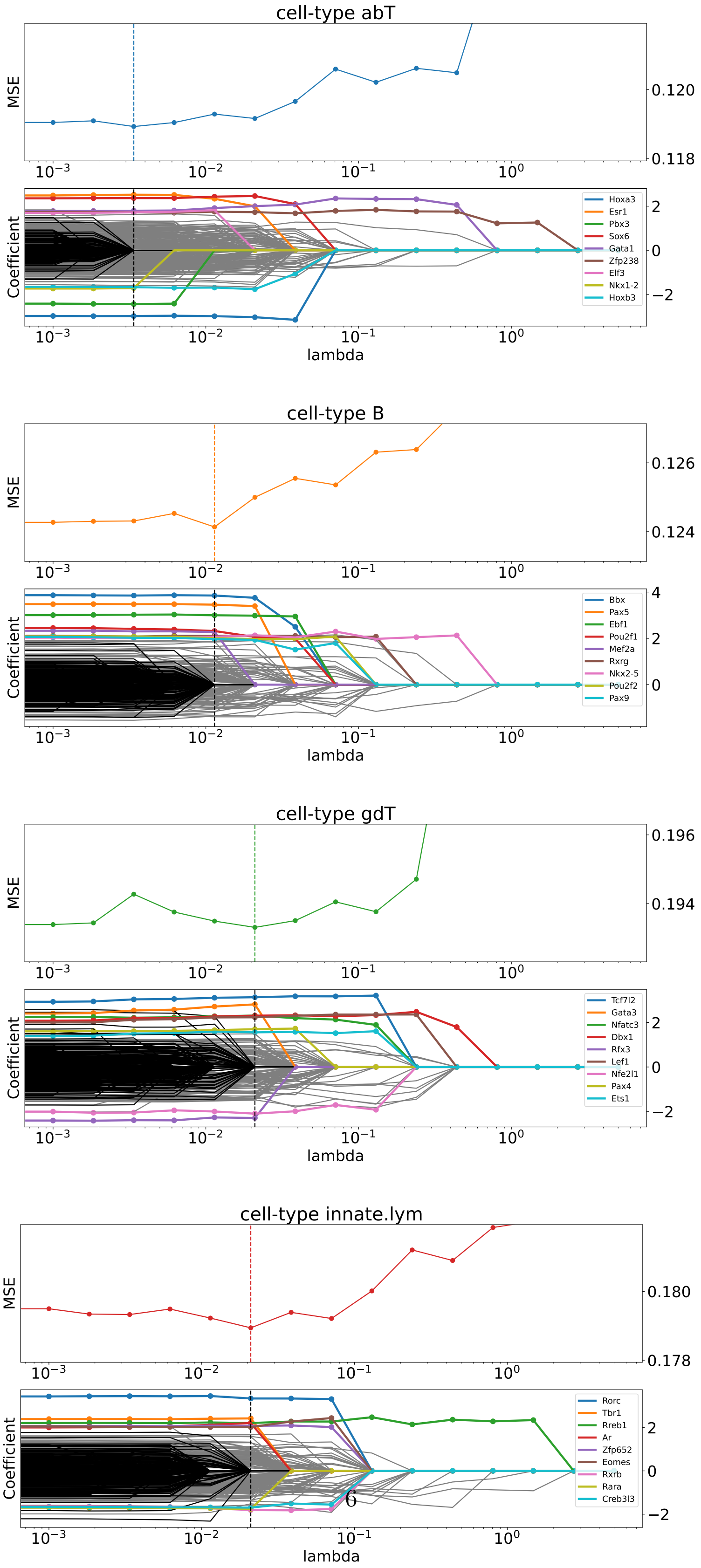


Figure S3: All cell-types and their corresponding PATH plots.

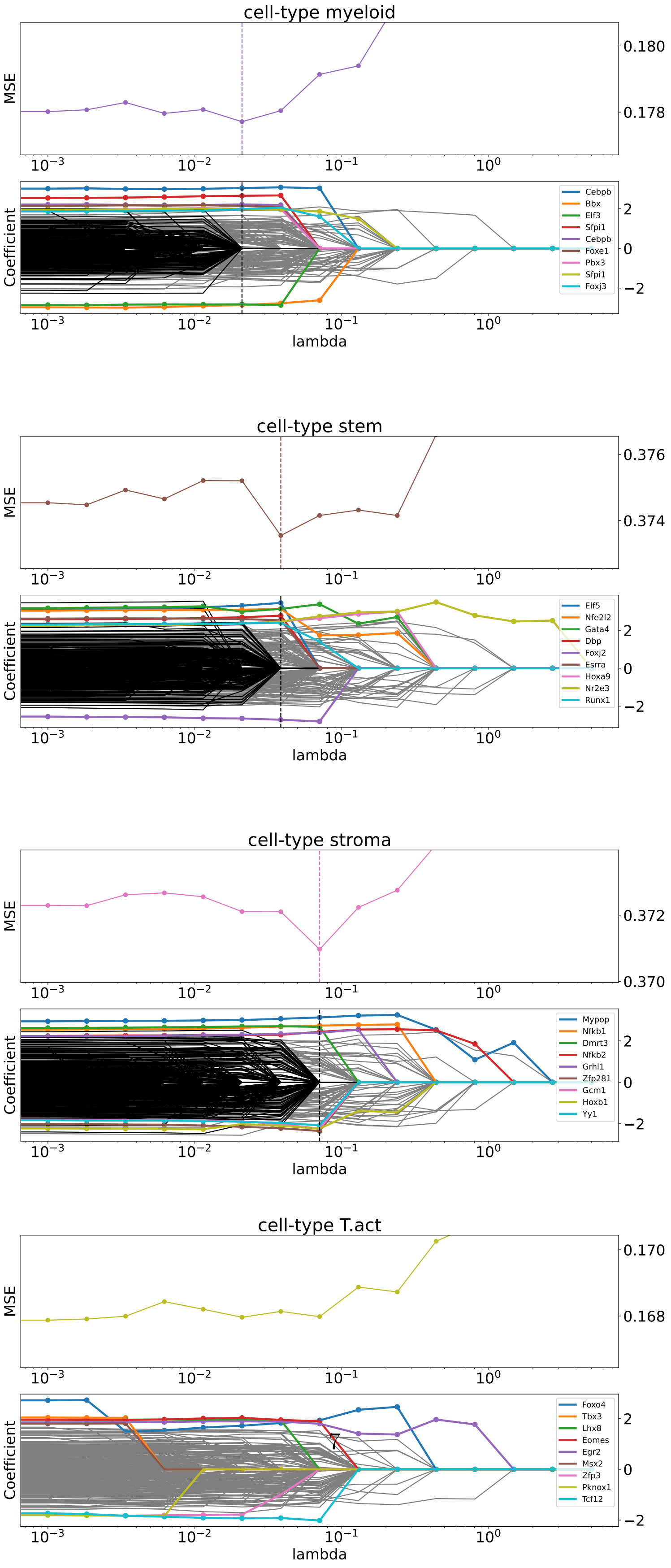


Figure S3: All cell-types and their corresponding PATH plots (cont.).

| transition | variance | MSE | R2 |
|---|---|---|---|
| LTHSC.34+.BM-LTHSC.34-.BM | 2.07 | 2.02 | 0.02 |
| STHSC.150-.BM-LTHSC.34+.BM | 1.95 | 1.91 | 0.02 |
| MMP3.48+.BM-STHSC.150-.BM | 1.82 | 1.81 | 0.01 |
| MMP4.135+.BM-STHSC.150-.BM | 1.47 | 1.43 | 0.02 |
| proB.CLP.BM-MMP4.135+.BM | 1.40 | 1.34 | 0.05 |
| proB.FrA.BM-proB.CLP.BM | 1.16 | 1.15 | 0.01 |
| proB.FrBC.BM-proB.FrA.BM | 2.10 | 1.86 | 0.11 |
| preB.FrD.BM-proB.FrBC.BM | 1.93 | 1.89 | 0.02 |
| B.FrE.BM-preB.FrD.BM | 1.97 | 1.94 | 0.02 |
| B1b.PC-B.FrE.BM | 2.43 | 2.37 | 0.02 |
| B.Sp-B.FrE.BM | 2.20 | 2.17 | 0.02 |
| B.Fem.Sp-B.FrE.BM | 2.43 | 2.34 | 0.04 |
| B.T1.Sp-B.FrE.BM | 1.86 | 1.84 | 0.01 |
| B.T2.Sp-B.T1.Sp | 2.02 | 1.98 | 0.02 |
| B.T3.Sp-B.T2.Sp | 2.05 | 2.04 | 0.00 |
| B.Fo.Sp-B.T3.Sp | 1.93 | 1.93 | -0.00 |
| B.MZ.Sp-B.FrE.BM | 2.32 | 2.27 | 0.02 |
| B.mem.Sp-B.Fo.Sp | 2.08 | 2.06 | 0.01 |
| B.GC.CC.Sp-B.Fo.Sp | 2.24 | 2.18 | 0.03 |
| B.GC.CB.Sp-B.GC.CC.Sp | 1.63 | 1.60 | 0.02 |
| B.PB.Sp-B.Fo.Sp | 2.73 | 2.62 | 0.04 |
| B.PC.Sp-B.PB.Sp | 2.33 | 2.27 | 0.03 |
| B.PC.BM-B.PC.Sp | 2.42 | 2.35 | 0.03 |
| preT.DN1.Th-MMP4.135+.BM | 1.34 | 1.29 | 0.03 |
| preT.DN2a.Th-preT.DN1.Th | 1.20 | 1.17 | 0.02 |
| preT.DN2b.Th-preT.DN2a.Th | 1.51 | 1.40 | 0.07 |
| preT.DN3.Th-preT.DN2b.Th | 1.56 | 1.54 | 0.01 |
| T.DN4.Th-preT.DN3.Th | 1.66 | 1.63 | 0.02 |
| T.ISP.Th-T.DN4.Th | 1.71 | 1.71 | 0.00 |
| T.DP.Th-T.ISP.Th | 2.01 | 1.97 | 0.02 |
| T.4.Th-T.DP.Th | 2.19 | 2.03 | 0.07 |
| T.8.Th-T.DP.Th | 2.26 | 2.06 | 0.09 |
| T.4.Nve.Sp-T.4.Th | 2.10 | 2.08 | 0.01 |
| Treg.4.25hi.Sp-T.4.Th | 2.35 | 2.28 | 0.03 |
| NKT.Sp-T.4.Th | 2.53 | 2.43 | 0.04 |
| Treg.4.FP3+.Nrplo.Co-T.4.Nve.Sp | 2.67 | 2.54 | 0.05 |
| T.4.Sp.aCD3+CD40.18hr-T.4.Nve.Sp | 2.18 | 2.11 | 0.03 |
| T.8.Nve.Sp-T.8.Th | 1.80 | 1.77 | 0.02 |
| T8.TN.P14.Sp-T.8.Nve.Sp | 1.70 | 1.68 | 0.01 |
| T8.TE.LCMV.d7.Sp-T.8.Nve.Sp | 2.35 | 2.26 | 0.04 |
| T8.MP.LCMV.d7.Sp-T.8.Nve.Sp | 2.21 | 2.13 | 0.03 |
| T8.IEL.LCMV.d7.Gut-T.8.Nve.Sp | 2.40 | 2.32 | 0.03 |
| T8.Tcm.LCMV.d180.Sp-T8.MP.LCMV.d7.Sp | 2.09 | 2.06 | 0.01 |
| T8.Tem.LCMV.d180.Sp-T8.MP.LCMV.d7.Sp | 2.20 | 2.19 | 0.00 |
| Tgd.g2+d17.24a+.Th-T.DN4.Th | 1.90 | 1.85 | 0.02 |
| Tgd.g2+d1.24a+.Th-T.DN4.Th | 2.01 | 1.96 | 0.02 |
| Tgd.g1.1+d1.24a+.Th-T.DN4.Th | 1.93 | 1.78 | 0.08 |
| Tgd.Sp-T.DN4.Th | 2.22 | 2.09 | 0.06 |
| Tgd.g2+d17.LN-Tgd.g2+d17.24a+.Th | 3.00 | 2.84 | 0.05 |
| Tgd.g2+d1.LN-Tgd.g2+d1.24a+.Th | 2.46 | 2.40 | 0.02 |
| Tgd.g1.1+d1.LN-Tgd.g1.1+d1.24a+.Th | 2.24 | 2.12 | 0.05 |
| NK.27+11b-.Sp-MMP4.135+.BM | 2.97 | 2.50 | 0.16 |
| NK.27+11b+.Sp-NK.27+11b-.Sp | 1.56 | 1.54 | 0.01 |
| NK.27-11b+.Sp-NK.27+11b+.Sp | 1.82 | 1.82 | -0.00 |
| NK.27+11b-.BM-MMP4.135+.BM | 2.85 | 2.38 | 0.16 |
| NK.27+11b+.BM-NK.27+11b-.BM | 1.47 | 1.47 | -0.00 |
| NK.27-11b+.BM-NK.27+11b+.BM | 1.83 | 1.79 | 0.02 |
| ILC2.SI-MMP4.135+.BM | 3.16 | 2.67 | 0.15 |
| ILC3.NKp46-CCR6-.SI-MMP4.135+.BM | 3.22 | 2.67 | 0.17 |
| ILC3.CCR6+.SI-MMP4.135+.BM | 3.14 | 2.60 | 0.17 |
| ILC3.NKp46+.SI-MMP4.135+.BM | 3.23 | 2.71 | 0.16 |
| GN.BM-MMP3.48+.BM | 3.52 | 3.22 | 0.09 |
| GN.Sp-GN.BM | 2.26 | 2.23 | 0.01 |
| GN.Thio.PC-GN.Sp | 2.06 | 2.04 | 0.01 |
| DC.103+11b+.SI-MMP3.48+.BM | 3.15 | 2.77 | 0.12 |
| DC.103+11b-.SI-MMP3.48+.BM | 3.15 | 2.70 | 0.14 |
| DC.8+.Sp-MMP3.48+.BM | 3.28 | 2.81 | 0.14 |
| DC.4+.Sp-MMP3.48+.BM | 2.99 | 2.68 | 0.10 |
| DC.pDC.Sp-MMP3.48+.BM | 2.98 | 2.68 | 0.10 |
| Mo.6C+II-.Bl-MMP3.48+.BM | 3.09 | 2.72 | 0.12 |
| Mo.6C-II-.Bl-MMP3.48+.BM | 3.26 | 2.91 | 0.11 |
| MF.226+II+480lo.PC-MMP3.48+.BM | 3.25 | 2.73 | 0.16 |
| MF.RP.Sp-MMP3.48+.BM | 3.35 | 2.97 | 0.12 |
| MF.PC-MMP3.48+.BM | 3.25 | 2.81 | 0.14 |
| MF.Fem.PC-MMP3.48+.BM | 3.41 | 2.99 | 0.12 |
| MF.SI-Mo.6C+II-.Bl | 2.12 | 2.03 | 0.04 |
| MF.microglia.CNS-Mo.6C+II-.Bl | 2.90 | 2.72 | 0.06 |
| MF.Alv.Lu-Mo.6C-II-.Bl | 2.35 | 2.27 | 0.03 |
| MF.pIC.Alv.Lu-MF.Alv.Lu | 1.54 | 1.53 | 0.01 |
| NKT.Sp.LPS.3hr-NKT.Sp | 2.12 | 2.11 | 0.00 |
| NKT.Sp.LPS.18hr-NKT.Sp.LPS.3hr | 1.99 | 1.97 | 0.01 |
| NKT.Sp.LPS.3d-NKT.Sp.LPS.18hr | 1.95 | 1.94 | 0.01 |

Table S2: The complete results for tree-diff dataset. The variance corresponds to the variance in the output while the R2 and MSE are a measure of prediction accuracy

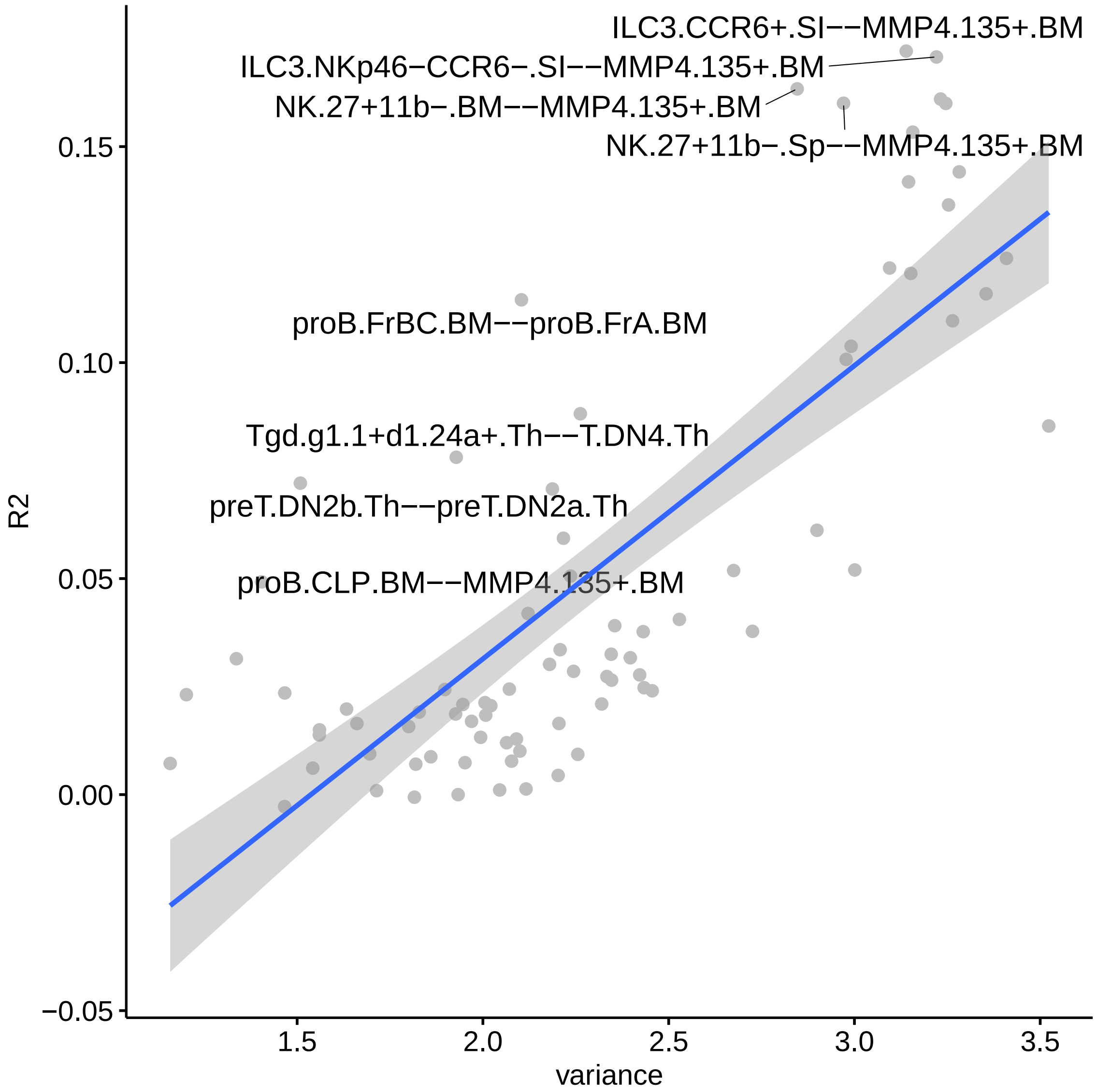


Figure S4: Relationship between the variance in OCR activity difference and its predictability as measured by $R^2$. Generally the two are highly correlated though some transitions are clear outliers.
